# Pin Electrode Reactor: A novel cold atmospheric plasma device and its potential in glioblastoma treatment

**DOI:** 10.1101/2021.01.08.425903

**Authors:** Andressa Maria Aguiar de Carvalho, Sean Behan, Laurence Scally, Chaitanya Sarangapani, Renee Malone, Patrick J. Cullen, Brijesh Tiwari, James F. Curtin

## Abstract

Glioblastoma multiforme (GBM) is the most common and biologically aggressive brain tumour. The current standard therapy for GBM consists in surgical resection, followed by radiotherapy and chemotherapy. Yet, the treatment is limited due to the area for the surgical resection and for the inability of some drugs to cross the brain blood barrier, leading to a general prognostic of no more than a year. Cold atmospheric plasma (CAP) is a new approach in the treatment of this challenging disease. CAP interaction with cells is dependent on physical and chemical factors, with different plasma discharges, cell type, and culture conditions leading to different CAP activity. Considering the plasma self-adaptation that different plasma discharge modes can undergo, which leads to different interaction plasma/cells, the characterization of a new device is essential. In this study we analysed the effect of a novel large pin-to-plate non-thermal atmospheric plasma on U-251 MG cells under different conditions. The analysis of reactive oxygen and nitrogen species (RONS) on plasma, media and cells were also assessed. We were able to demonstrate that the pin-to-plate device is cytotoxic to GBM cells in a dose, time and ROS dependent manner. The measurements of RONS on plasma/media also give us an insight on the chemical effect of this novelty device, and the possibility to better understand the use of this device as a promising GBM therapy.

## Introduction

Glioblastoma multiforme (GBM) is a grade IV cancer, being the most malignant and frequent tumour of the brain (Hayat, 2012; Khani *et al.*, 2019). It has an average annual incidence of 11.833 per 100.000 population (Ostrom *et al.*, 2019), with a five-year relative survival of only 6.8 % (Hanif *et al.*, 2017; Ostrom *et al.*, 2019). Currently, GBM therapy consists of surgical resection, followed by radiation therapy and use of chemotherapy (Stupp *et al.*, 2005; Tamimi and Juweid, 2017). However, despite advances in science, the overall survival rate for GBM has not improved over the years (GLOBOCAN, 2018). It is still largely incurable, with a life expectancy of 14 months following diagnosis (Hanif *et al.*, 2017). These statistics highlight the need for new therapeutic approaches to overcome this challenging disease.

Plasma is described as the fourth state of matter and can be conceptualized as a neutral ionized gas composed of charged and neutral particles, as well as electrons, radicals, UV-radiation and electric fields (Yan, Sherman and Keidar, 2017; Semmler *et al.*, 2020). Cold atmospheric plasma (CAP) is generated at conditions of near atmospheric pressures, and at temperatures less than 40 *°C* (von Woedtke *et al.*, 2013; Conway *et al.*, 2016; Yan, Sherman and Keidar, 2017), which facilitates its application in the bio-medical field. Due to its unique characteristics CAP has been used in a wide range of biomedical applications, more recently it has become a promising anti-cancer modality, with studies demonstrating its cytotoxicity effects, selectivity potential and efficacy as a GBM therapy (Vermeylen *et al.*, 2016; Yang *et al.*, 2018; Privat-Maldonado *et al.*, 2018).

Different plasma discharge modes can lead to different interactions between plasma and cells, a phenomenon that Keidar *et al.*, (2018) defined as plasma self-adaptation. These changes are modulated through differences in the power supply, the gas composition and the distance between the plasma source and the cells (Keidar *et al.*, 2018). Cell type, cancer type, and culture conditions can also strongly influence CAP treatment (Biscop *et al.*, 2019; Semmler *et al.*, 2020). The interaction between CAP and mammalian cells depends on many physical and chemical factors. Previous studies have reported that physical factors including UV, heat as well as electromagnetic fields are negligible contributors to DNA damage (Kalghatgi *et al.*, 2011; Yan, Sherman and Keidar, 2017). In contrast, chemical factors have been observed to be responsible for the cellular response induced by CAP treatment, mainly due to its production of reactive oxygen species (ROS) (Graves, 2012; Yan, Sherman and Keidar, 2017). Reactive species generated in the gas phase are then transferred to the liquid medium. This represents a unique scenario based on variable life-times, free diffusion path lengths and multiple potentials of interactions (Bauer *et al.*, 2019). Plasma generates long and short-lived reactive species (Tanaka *et al.*, 2018). Amongst those species, oxygen (O_2_•, O, and O_3_, OH_•_, H_2_O_2_, HO_2_) and nitrogen species (N_2_ +, NO) play a key role in CAP induced effects (Conway *et al.*, 2016; Tanaka *et al.*, 2018; Semmler *et al.*, 2020).

The design of the device itself along with its input power is another characteristic to consider in the evaluation of biological activity. A pin-to-plate based electrode is one of the designs that have been developed (Zhang *et al.* 2014). The array of pins and a plate electrode that act as the ground provides lower values of power consumption and a stable and more diffusive nature of the discharge over the surface. The discharge was also reported to diffuse to a much larger area than the pin tip. Those characteristics would create an environment that allows for more interaction between reactive species and cells, working in an unmodified atmosphere (Liu *et al.*, 2013; Zhang *et al.*, 2014).

In the current study, the *in vitro* cytotoxicity of pin-to-plate on U-251 MG GBM cells were analysed under different conditions, in order to establish the optimum setting of this novel device and to understand its cytotoxicity potential. Antioxidants were also used to elucidate the role of RONS in the observed cytotoxicity. The presence of RONS was analysed on the discharge plume, media and cells, in order to gain a better understanding of the role of reactive species involved in the cytotoxicity process. This work provides a preliminary study into cytotoxicity and reactive species present on the pin-to-plate system, demonstrating the potential of this device in the treatment of GBM.

## Materials and Methods

### Chemicals

All chemicals used in this study were supplied by Sigma-Aldrich - Merck Group (Arklow, Ireland), unless stated otherwise.

### Cell Culture

The Human glioblastoma multiforme cell line (U-251MG, formerly known as U-373 MG-CD14), was a gift from Michael Carty (Trinity College Dublin). Cells were maintained in - Dulbecco’s Modified Eagle Medium (DMEM) supplemented with 10 % FBS - Fetal bovine serum and 1 % penicillin/streptomycin. Cells were routinely subcultured when 80% confluence was reached using 0.25 % w/v Trypsin solution. Cells were maintained in a humidified incubator at 37 °C containing 5 % CO_2_.

### Pin-to-plate device

This device is a novel large pin-to-plate non-thermal atmospheric plasma (Figure 1 A and B). It consists of an 88 pin stainless steel electrode, paired with a flat stainless steel ground plate powered by an AC Supply (Leap100, PlasmaLeap Technologies, Dublin, Ireland) that has an electrode matched resonance frequency of 30 - 125 kHz, a discharge frequency of 50 Hz - 3000 Hz with the power range from 50 W to 400 W and a discharge gap of a maximum of 55 mm. The pins are arranged in a slightly convex manner, with pins being closest to the ground plate at the centre at a distance of 40mm with a gradual increase to 55 mm at the outer edge. The air gap between the pin electrode and the ground plate serves as the sample treatment area, with all samples in the current study being placed in the centre. The samples were treated at a resonant frequency of 55.51 kHz, with a discharge frequency of 1000 Hz and duty cycles of 54 μs and 73 μs. The discharge gap was kept at 40 mm. Cells were treated for a time range of 0 - 320 seconds.

**Figure 1.**
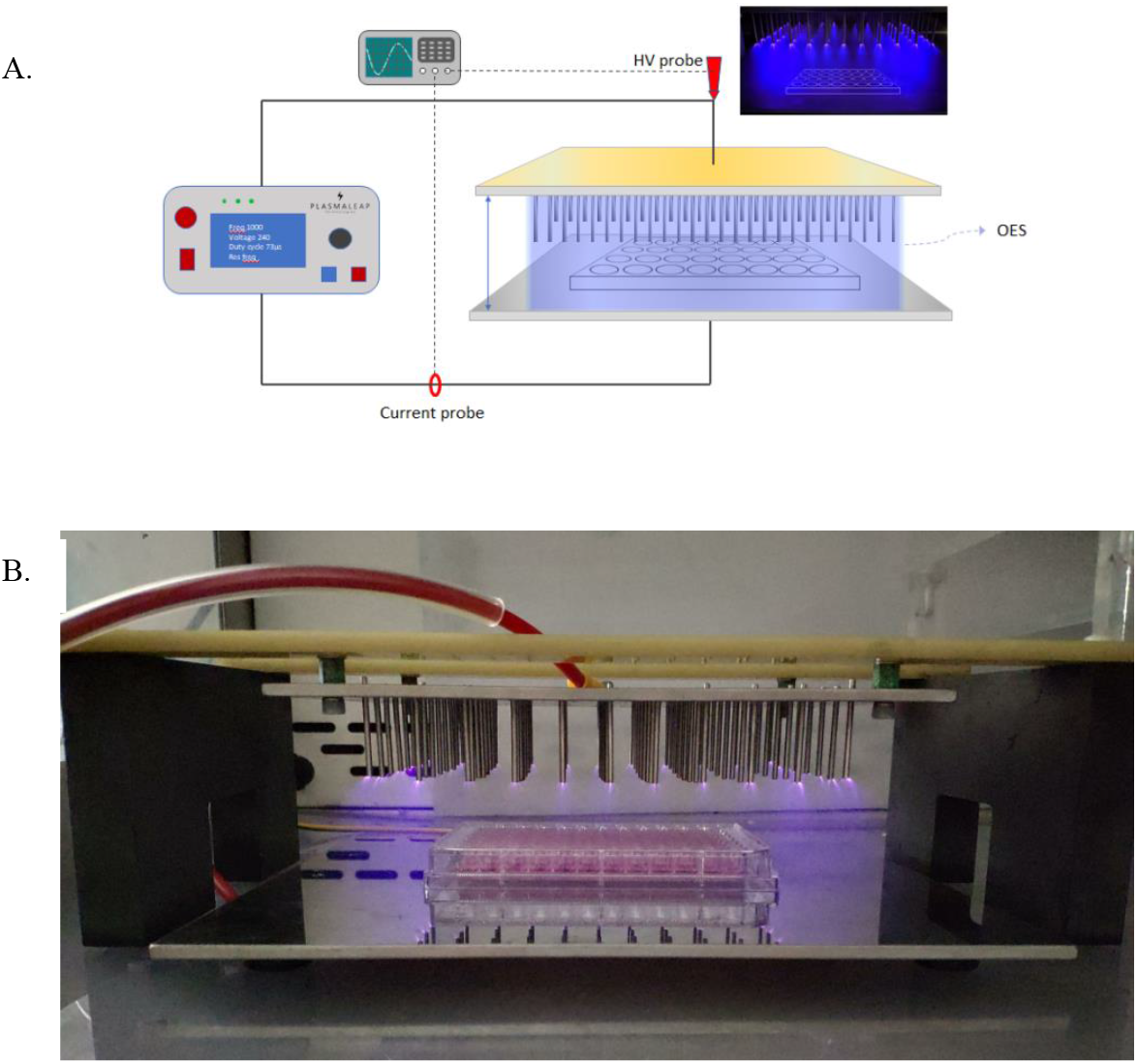
Pin-to-plate device. A - Schematic of pin-to-plate device B - Image of the pin-to-plate device demonstrating the position of the microplates for cell treatment.

### Cell viability assay

Cells were seeded on flat bottom 96 well-plates in 100 μL DMEM (Sarstedt, Ltd., Wexford, Ireland) and left to adhere overnight at 37 °C in a humidified atmosphere at a density of 2 x 10^3^ cells/well or 1 x 10^4^ cells/well for 96 hours incubation time post treatment and 24 hours post treatment, respectively. DMEM media without sodium pyruvate, and DMEM media supplemented with 0.11 g/L sodium pyruvate were used in this study where indicated. The 24 central wells in the 96 well plates were seeded, to ensure symmetry in plate loading to help stabilise the plasma field during treatment. The following day, 80 μL of media was removed, leaving 20 μL of media for treatment in each well, unless otherwise specified. Cells were then treated with pin-to-plate discharge at seven different time points (5, 10, 20, 40, 80, 160 and 320 seconds) using a voltage of 240 V, frequency of 1000 Hz and using a duty cycle of 54 μs or 73 μs. 80 μL of fresh media was added after treatment and cells were incubated at 37 °C using 5 % CO_2_ for 96 hours or 24 hours. Dimethyl sulfoxide (DMSO) (20 %) was used as positive control.

Cell viability was analysed using Alamar Blue™ Cell Viability Reagent (Thermo Fisher Scientific), an oxidation-reduction indicator that undergoes colorimetric and fluorometric change in response to cellular metabolic reduction. Cells were washed with sterile phosphate buffered saline (PBS) and incubated for 3 hours at 37 °C with a 10 % Alamar Blue™ solution in DMEM without FBS. Fluorescence was measured using an excitation wavelength of 530 nm and an emission wavelength of 595 nm on Varioskan Lux multi-plate reader (Thermo Scientific).

All experiments were performed at least three independent times with a minimum of 24 replicates per experiment.

### Live/Dead Cell Staining Using Propidium Iodide (PI)

Cells were seeded in 35 mm cell culture dishes (Thermo Fisher Scientific) to achieve a density of 5 x 10^5^ cells per culture dish on the day of the assay. Prior to CAP treatment culture media was aspirated from the dishes and replaced with 200 μL of DMEM media. The cells were then treated using the pin-to-plate system and immediately replenished with 2 mL of media. The cells were then incubated for their allotted time, 4 or 96 hours.

For PI staining the media was removed and the cells washed with PBS and harvested using Trypsin. The removed media, PBS wash and harvested cells were retained and combined into a single tube for centrifugation at 250 x g for 5 minutes. The supernatant was then aspirated, and the pellet resuspended in 1 mL of PBS. PI was then added to the cell suspension at 10 μg/mL and incubated for 1 minute. PI fluorescence was then measured with the BD Accuri C6 Plus flow cytometer (BD, Oxford, UK) using the FL2 channel.

### Optical measurement

In order to obtain preliminary understanding and to quantify the gas chemistry of the plasma discharge and the interactions that may occur at the sample boundary of the cell culture media during treatment, the use of optical emission spectroscopy (OES) and optical absorption spectroscopy (OAS) was employed.

20 μL of DMEM media was added into a 96 well-plate. Following positioning the plate at the center of the device, both OES and OAS were carried out using an Edmund Optics CCD spectrometer (Model: BRC115U) with a wavelength range of 200 – 850 nm. Software BWSpecTM V4.02 was used to record the spectra for both OES and OAS during the measurement processes. The spectral resolution for the spectrometer used in the current study, was between 0.6 and 1.8 nm and is wavelength dependent. OES and OAS readings were taken across the top (1.3 cm above the ground plate) and at the centre of the 96 well-plate and then 1.5 cm to the left and right of that.

OES: The acquisition of each spectrum was carried out with an integration time of 2000 ms and the time between each spectrum was 5.5 seconds. In order to carry out these measurements, a fibre optic cable was used that had an adjustable lens attached to the end and was directed perpendicular to the system and across the top of the well-plate. Following OES measurements, the spectra were analysed by integrating the area under each peak of interest to calculate the value of the total intensity in arbitrary units. Further analysis, through the use of equation (1), gives an insight into the electron energy distribution. By taking the line intensity ratio of N2(337nm)/N2^+^(391nm), the ratio of low energy electrons to high energy electrons can be found. This gives a better understanding of what reaction mechanisms may occur and which excitation routes are most likely for the generation of reactive species within the plasma discharge (Scally *et al.*, 2018).

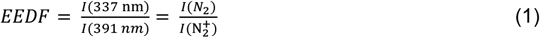

In order to carry out the OAS measurement of the generated plasma, two fibre optic cables were used, each of which had adjustable lenses attached to one end to optimise the focal point of detection. They were placed perpendicular to one another at a distance of 25 cm and allowed to have an unobstructed direct line-of-sight between them. One fibre optic cable was connected to a deuterium-tungsten UV-Vis-NIR light source and the other was connected to the CCD spectrometer to detect the incoming light. By referencing the incoming light to the incoming light during plasma discharge, the average optical density of different absorbing species can be determined. The spectra were measured with an integration time of 400 ms with 1600 ms set between each measurement. The spectra from the OAS measurements were analysed by using equation 2 to find the average spatial density of O_3_ during plasma discharge and post-discharge. In equation 2 *D*(*t*) is the average spatial density (cm^−3^), *L* is the optical path (cm), *I*(*0*) is the reference intensity with no plasma discharge (A.U.), *I*(*t*) is the measured intensity (A.U.) during and after plasma discharge, and *σ*(*λ*) is the wavelength dependent absorption cross-section for the species of interest. For O_3_, the wavelength of optimal absorption is taken as 253.7 nm which gives a cross-section value of 1.154 x 10^−17^ cm^2^.

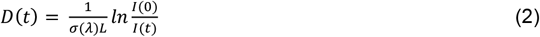

### Chemical Analysis of Reactive Species in Cell Culture Medium

Nitrite and hydrogen peroxide concentrations were quantified using Griess reagent for nitrite and Amplex™ Red Hydrogen Peroxide/Peroxidase Assay Kit (ThermoFisher, Oregon, USA), respectively. The assays were performed in a flat bottom 96-well plate (Sarstedt, Ltd., Wexford, Ireland). 20μl of DMEM in the absence of sodium pyruvate and phenol red were exposed to pin-to-plate discharge at four different time points (5, 20, 80 and 160 seconds), 240 V, 1000 Hz with a variance in duty cycle (54 μs and 73 μs). Plates were incubated for 24 hours before concentrations were quantified as described.

For nitrite quantification, 50 μL of Griess reagent for nitrite was added to 50 μL of media (20 μL of treated media + 30 μL of fresh media) and incubated at room temperature for 30 minutes in the dark. Absorbance was read at 548 nm. Hydrogen peroxide was quantified according to the kit protocol. 50 μl of a working solution of 100 uM Amplex™ red reagent and 0.2 U/mL Horse Radish Peroxidase (HRP) were added to 50 μL of media (20 μL of treated media + 30 μL of fresh media). Following 30 mins incubation at room temperature, in the dark, fluorescence was measured using an excitation wavelength of 530 nm and emission of 590 nm. A standard curve of sodium nitrite (0 – 100 μM) and hydrogen peroxide (0 - 20μM) was used.to determine nitrite and hydrogen peroxide concentrations.

All experiments were performed in triplicates.

### Analysis of Reactive Species in the U-251MG cell line

Reactive species in U-251MG cells were quantified by flow cytometry (CytoFlex, Beckman Coulter) using the cell-permeant 2’,7’ dichlorodihydrofluorescein diacetate (H2DCFDA) dye, a chemically reduced form of fluorescein that after diffusion into the cells is deacetylated and later oxidized by ROS into highly fluorescent 2’,7’-dichlorofluorescein (DCF). Cells were seeded at a density of 2.5 - 3 x 10^5^ into TC Dish 60 (Sarstedt, Ltd, Germany) using DMEM high glucose in the absence of sodium pyruvate and left to adhere overnight. Prior to treatment, media was removed and cells washed with sterile PBS. Cells were incubated for 1 hour at 37 *°C* with 25 μM of H2DCFDA using DMEM in the absence sodium pyruvate and phenol red. 2.7 mL of media was then removed and the remaining 0.3 mL of media was treated. Cells were exposed to pin-to-plate discharge at 3 different time points (20, 80 and 160 seconds), at 240 V, 1000 Hz and using either 54 μs or 73 μs as duty cycle. After 3 hours of incubation at 37 *°C* using 5 % CO_2_, cells were collected and quantified using FL-1 green channel. The experiments were replicated twice.

### ROS scavenger assays

Reactive oxygen species inhibitor N-acetyl cysteine (NAC) was used as ROS scavenger. Cells were seeded as previously described and left to adhere overnight. For dose response curve, cells were incubated for 1 hour at 37 °C with 4 mM NAC in DMEM in the absence of pyruvate. 80 μL of this solution was removed and cells were then exposure to pin-to-plate discharge. After treatment fresh media containing 4 mM of NAC was added and cells were incubated at 37 *°C* using 5 % CO_2_ for 96 hours. Cell viability was measured using the Alamar Blue™assay. NAC titration was performed exposing cells to a range of 0 - 8 mM, cell viability was assessed after 96 hours.

### Statistical Analysis

All experiments were replicated at least 3 times unless otherwise stated. Dose response curve was measured using non-linear regression. Data is presented as mean ± SEM, multiple comparison analysis were performed using Sidak’s test, unless otherwise stated. Statistical analyses and curve fitting were performed using Prism 7, GraphPad Software, Inc. (USA). For ROS quantification, data was analysed using linear regression (Prism 7) interloping unknowns from standard curves. The values of the given concentrations were then plotted using Two-way ANOVA. CytExpert software was used for flow cytometry analysis and mean of FITC-A was use to plot the reading results in columns statistics. PI uptake studies were analysed using Two Way ANOVA with Tukey’s post-test.

## Results

### Pin-to-plate presents cytotoxicity towards GBM cells in a time/dose-dependent manner

Several studies have demonstrated the cytotoxicity and selective effects of cold atmospheric plasma on cancer cells (Vermeylen *et al.*, 2016; Yang *et al.*, 2017; Privat-Maldonado *et al.*, 2018). However, the effects of pin-to-plate devices on cancer cells have not yet been fully explored. Therefore the cytotoxicity of this novel plasma device towards U-251 MG was evaluated. Dose response curves were established by exposing cells to seven different doses of CAP (5 – 320 seconds) at 240 V, 73 μs and at 1000 Hz under different media conditions. Cell viability was then measured using the Alamar Blue™ assay.

Firstly, the effect of plasma discharge on GBM cells after 24 hours incubation time under two different media (DMEM high glucose with and without the presence of sodium pyruvate) was analysed. An IC50 of 143.6 seconds (116.5 s ± 176.9 s) and 145.8 seconds (119.1 s ± 178.6 s) was found for cells treated with DMEM without and with sodium pyruvate, respectively (Figure 2 A). Two-way ANOVA demonstrated that there is no significant difference in viability between the media used, however, doses of CAP present significant differences (P < 0.0001). Considering that after 24 hours post treatment the plasma discharge did not promote full cytotoxicity in the cells, even at the highest dose (320 seconds, cell viability = 37.23 - 39.43 %) it was decided to incubate the cells for a longer time after treatment. U-251 MG cells were therefore incubated for 96 hours post – treatment using the same conditions as the one described above (Figure 2 B). It was found that a longer incubation time appears to be needed for pin-to-plate doses to achieve their full cytotoxicity potential (320 seconds, cell viability = 6.26 – 0.01 %). When cells were incubated for 96 hours post-treatment, an IC50 of 5.716 seconds (5.308 s ± 6.155 s) for cells with DMEM in the absence of sodium pyruvate and an IC50 of 36.38 seconds (33.51 s ± 39.48 s) for cells with DMEM in the presence of sodium pyruvate was observed. There was a significant difference between IC50 DMEM used (P = 0.0001) in cells treated and incubated for 96 hours, which would suggest a protective effect of sodium pyruvate. In almost all multiple comparisons analysed, doses of CAP were significantly different (P < 0.0001). A full description of Sidak’s multiple comparisons test can been see in the supplementary Table I and II. The electrodes/sample treatment area is housed in a fitted container to minimize escape of CAP generated reactive species into the general environment. In order to define the effect that the containment would have on the cytotoxicity, cells were treated in the absence of the container box and incubated for 96 hours. Figure 2 B shows that cell viability is dependent on containment of the plasma during treatment with containment, with no cytotoxicity being observed for cells treated in the absence of the container.

**Figure 2.**
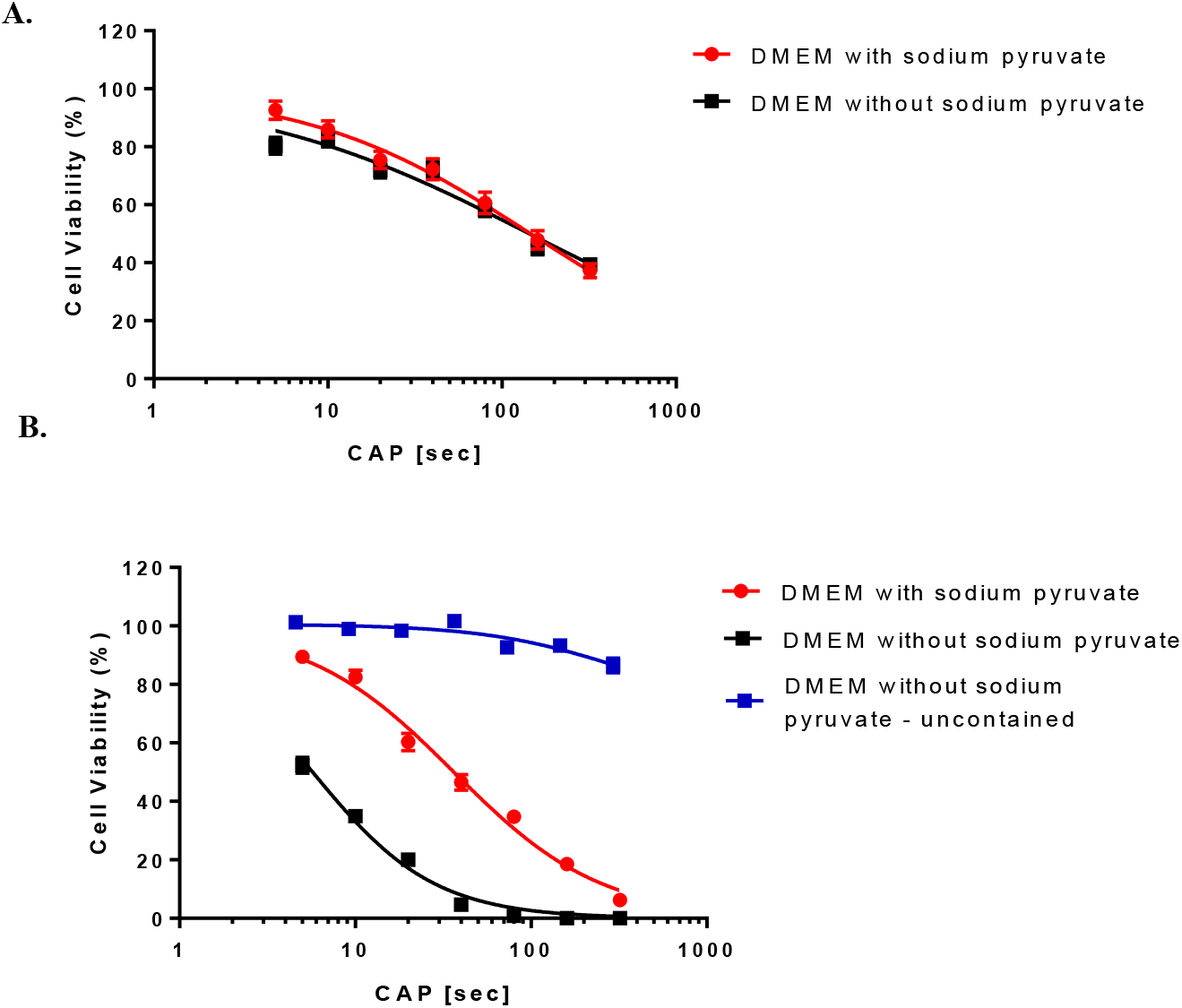
Pin-to-plate cytotoxicity in U-251 MG cells. To evaluated pin-to-plate cytotoxicity towards GBM cells and whether sodium pyruvate would have any protective effect, cells were exposed to plasma treatment at 240 V, 1000 Hz and 73 μs. Samples were treated at seven different doses of CAP and incubated for 24 hours post treatment (A) or 96 hours (B). To analyse the influence of plasma field containment, U-251 MG cells were treated without the containment box, using the same settings above and incubated for 96 hours (B).

From the experiments above it was established that cells would be incubated 96 hours post treatment for further experiments (when cytotoxicity or effects or its cytotoxicity want to be analysed). Media without sodium pyruvate would be used whenever the full effect of pin-to-plate discharge want to be evaluated. Despite the difference in IC50, all cells exposure to plasma discharge was susceptive in a dose depend manner. A table with all IC50, Hill Slope and coefficient of determination (R^2^) values of the dose response curve can been found in the supplementary table I and II.

Propidium iodide (PI) was used to secondarily confirm pin-to-plate induced cancer cell death, and to investigate its immediate toxicity. The cell membrane is impermeable to PI in normal conditions, so if PI DNA binding is observed in cells it indicates membrane damage/cell death or membrane permeabilization. PI uptake was measured 4 hours post treatment, with an increase of 4.8 % observed between the untreated control and 160 seconds of CAP treatment as seen in Figure 3. This contrasts to the 72.3 % increase of uptake observed in the 96 hours incubated samples. This corroborates the Alamar blue™ assay data (Figure 2) and shows that very little measurable membrane damage occurs immediately post treatment suggesting that cell death is programmed.

**Figure 3.**
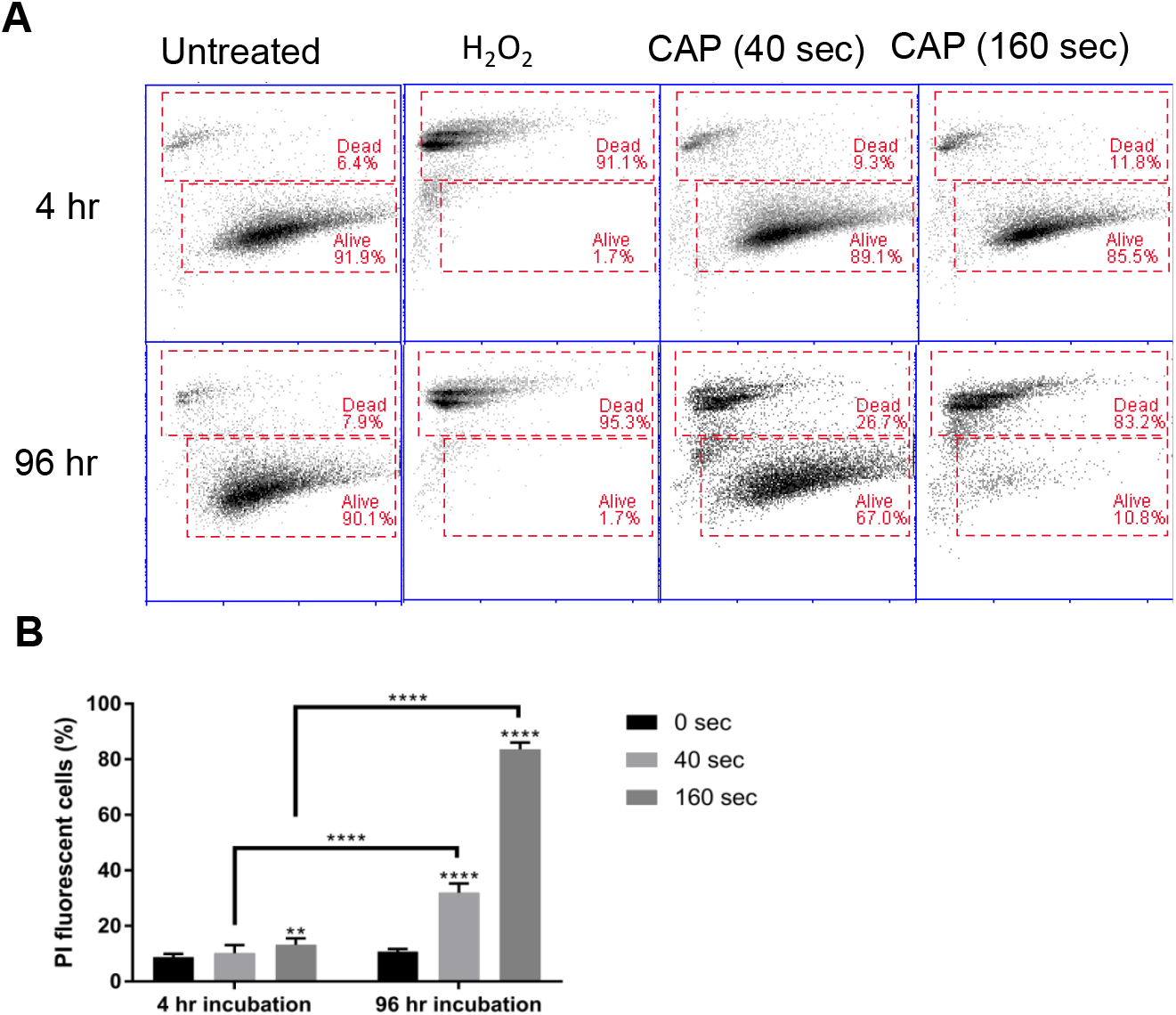
PI uptake in pin-to-plate treated U-251 MG cells. PI uptake was measured by flow cytometry and used as an indicator of cell death. Cells were treated at 240 V, 1000 Hz and 73 μs for either 40 or 160 seconds. PI uptake was then measured both 4 and 96 hours post treatment (A) and represented as a bar chart (B).

### Pin-to-plate presents RONS dependent cytotoxicity

The different responses observed at 1000 Hz in the presence and absence of sodium pyruvate may indicate the presence of ROS cytotoxic effects. The effect of duty cycle on ROS-induced cytotoxicity was therefore evaluated at 1000 Hz. NAC, a ROS scavenger, was then used. Figure 4 A and C demonstrated that NAC reversed the cytotoxicity effects of pin-to-plate device, with no significant difference (P = 0.7461) seen in the cytotoxicity between duty cycles. The highest protection was observed at 80 seconds of CAP treatment for both 73 μs and 54 μs, with a mean variation in IC50 of more than 150 fold for 73 μs (mean CAP only = 6.583 s; mean CAP/NAC = 1018 s).

**Figure 4.**
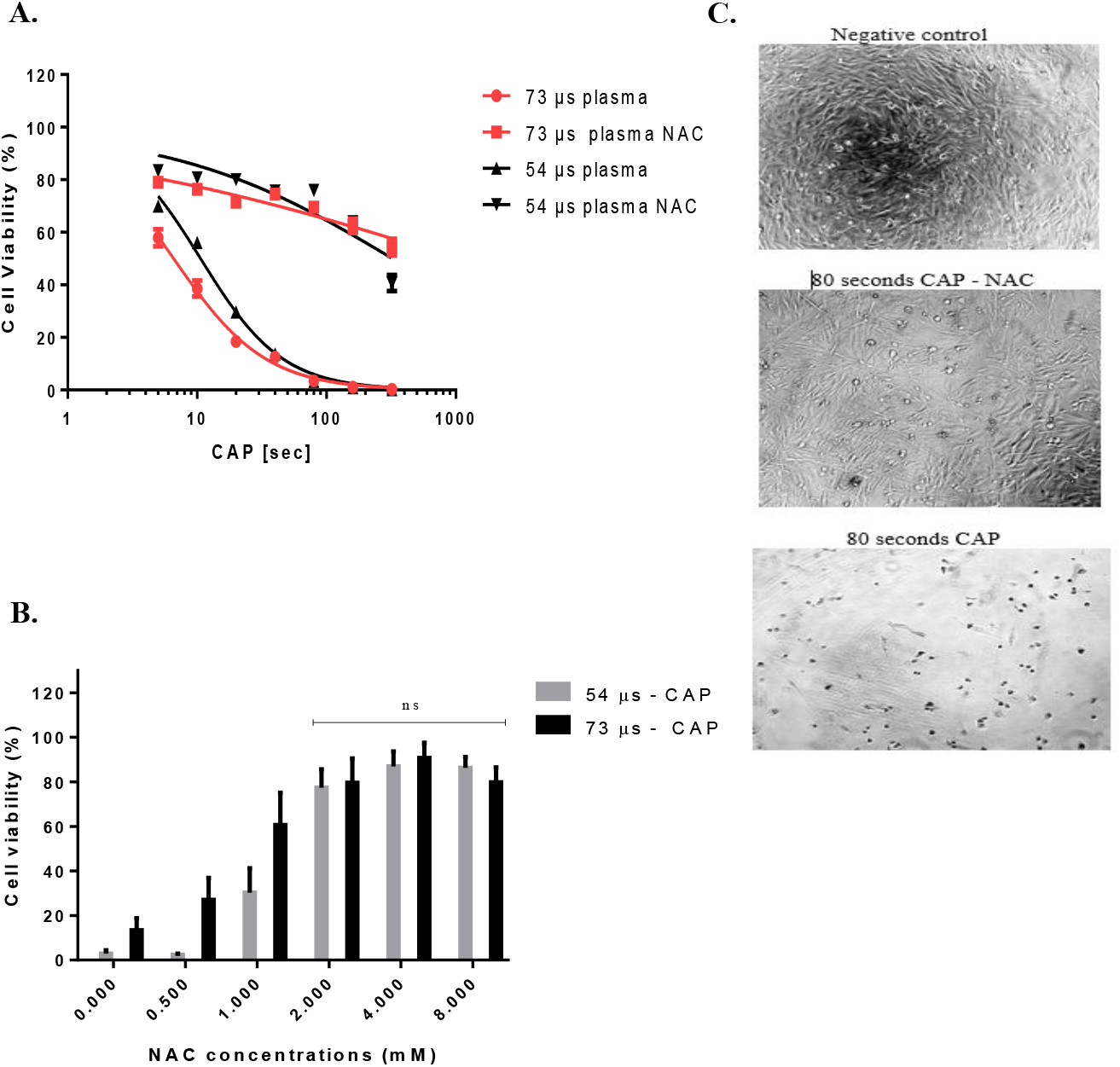
N-acetyl-L-cysteine as plasma cytotoxicity inhibitor. Cells were exposure to either NAC in DMEM without sodium pyruvate or DMEM only. After 96 hours incubation, cell viability was measured. (A) Dose response curve of NAC and plasma were evaluated for seven doses of CAP using non – linear regression. Unpaired t-test was also applied. (B) Titration of NAC was also analysed. Data shown as row statistic of normalized data, two-way ANOVA with Sidak’s multiple comparison was also performed. (C) Images of the protected effect of NAC were taken using optical microscope, brightness/contrast of images were automatically adjusted using imageJ. All experiments were performed in triplicate with a minimum of 12 replicates per experiment.

NAC significantly protected cells from cytotoxicity at each dose of CAP and duty cycle tested (P < 0.0001). Titration was performed to confirm the optimum working concentration. Figure 4 B demonstrates that there is no significant difference between 2 mM, 4 mM and 8 mM of NAC, showing that a 2-fold increase or decrease in NAC concentration does not change the protective effect.

### Spectrometer analysis of pin-to-plate plasma discharge

The *in vitro* anti-cancer effect of CAP relies on the interaction between CAP and cells (Yan, Sherman and Keidar, 2017). Therefore, the characterization of RONS generated on the discharge plume is one of the key elements in understanding CAP bioeffects. OES and OAS techniques, while simple in their employment, are powerful methods of determining the formation and stability of different reactive species within a plasma discharge. These analyses were carried out whilst simulating the same conditions applied for cytotoxicity assays.

Plasma discharge was analysed under 1000 Hz at two different duty cycles (54 μs and 73 μs) to elucidate whether any difference on the RONS production would been seen on the plasma discharge at those two different duty cycles.

For OES measurements, there was no detection of NO or O in the emission spectra obtained supplementary figure I), indicating that this device produces little to no species in the range were RONS emission exists. Following the initial spectroscopic measurement, it can be seen that the main reactive species that could be measured were OH (300 nm), N2 from the second positive system (SPS) (315, 337, 357, and 377 nm), and N2^+^ from the first negative system (FNS) (391 nm). From this, the temporal and spatial characterisation of the system was carried out using OES to quantify the energy distributions throughout the system, the stability of the discharge, and the possible mechanistic routes that occur within the discharge and at the sample boundary. From Figure 5, it is seen that there are fluctuations for all species over time. The analysis location on the plate also shows its variance. The right of the center, at 73 μs, presents the highest intensity of the species analysed. However, the calculated EEDF from the line ratio shows it highest intensity at the right center at 54 μs. Overall, OES analysis at 73 μs, shows an indication of higher intensity, although minor and arbitrary, when compared to 54 μs.

**Figure 5.**
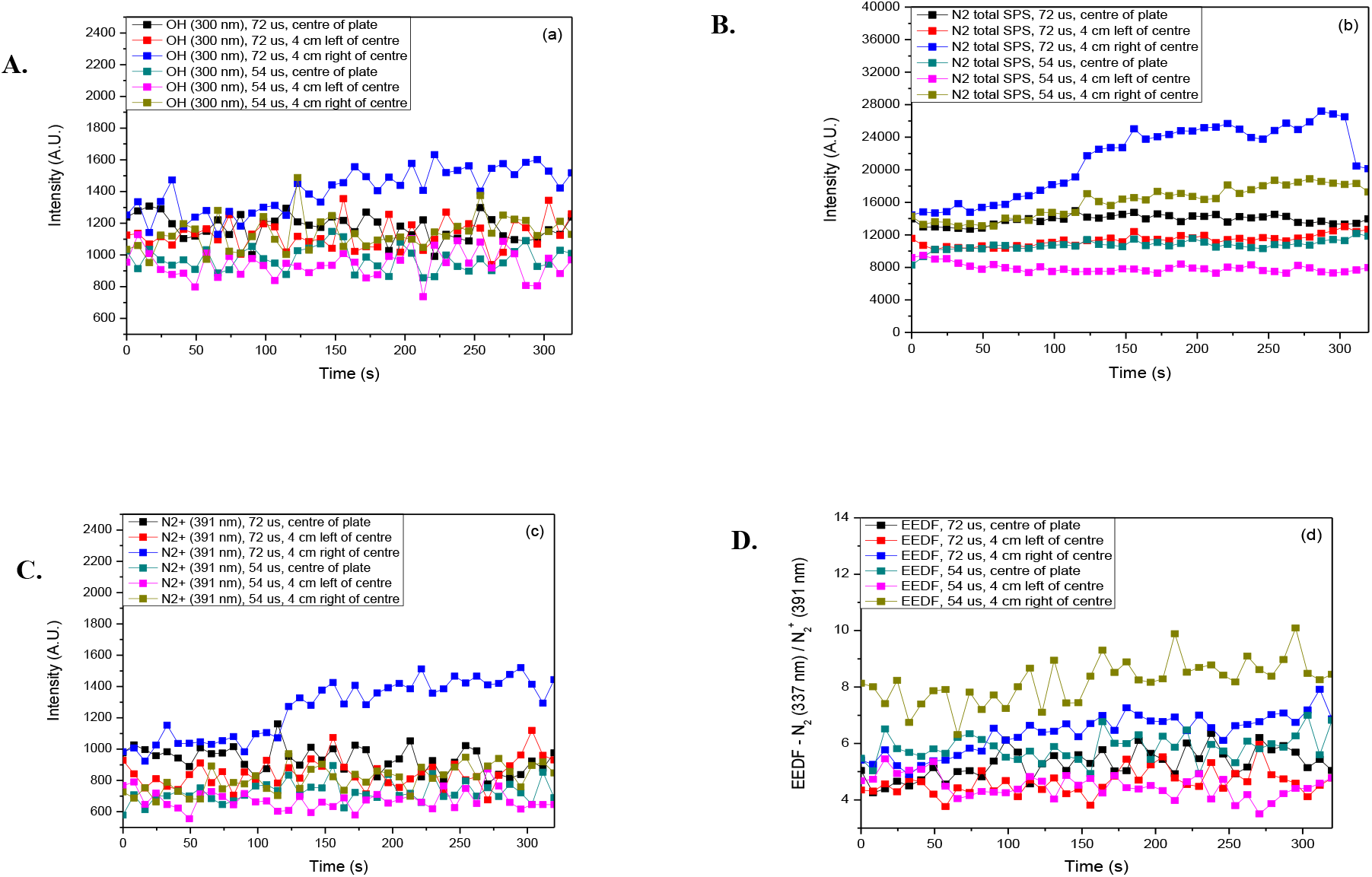
Spectroscopic data obtained with a 96-well plate placed into the central portion of the system for treatment. All graphs show the temporal and spatial evolution of species detected through OES and the changes for the energy of the system at 240 V, 1000 Hz and either 73 μs or 54 μs. (A) Changes observed for OH. (B) Sum total of measured emissions from the SPS of N2. (C) Evolution measured for the only detectable line from the FNS (N2+ 391 nm). (D). Calculated EEDF from the line ratio shown in equation 1 (see methodology).

OAS measurement demonstrated that most levels of O_3_ are relatively close and comparable to each other (Fig. 6). The two outliners (72 μs center of the plate and 72 μs 4 cm right from the center) are from the same position and settings that yielded higher levels of N2, N2^+^, and OH. It is also seen that O_3_ levels drop immediately when the device is turned off.

**Figure 6.**
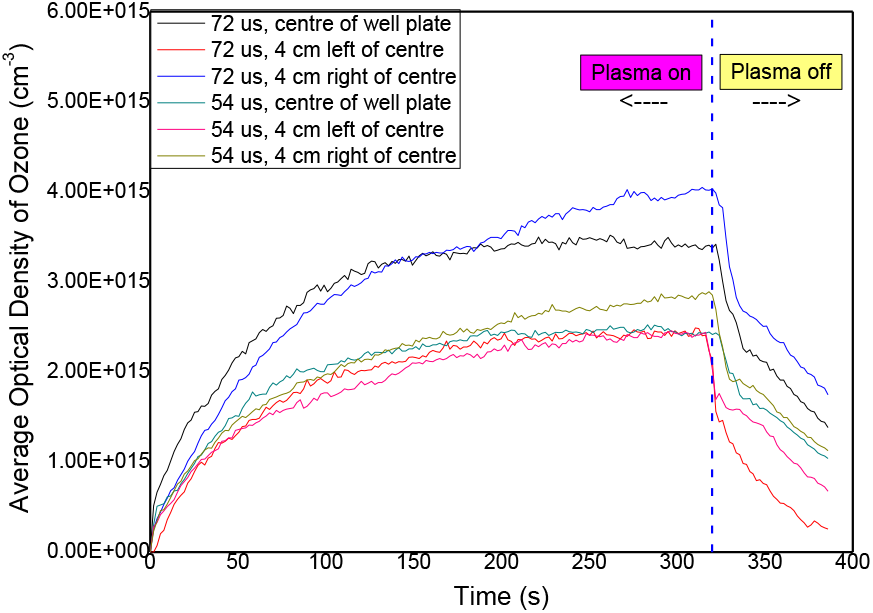
OAS measurement of O_3_ during plasma discharge and post-discharge. Analysis were performed at 240 V, 1000 Hz and either 73 μs or 54 μs.

### Quantification of nitrite and hydrogen peroxide in media

The quantification of these two species were performed, in order to gain a clearer understanding of RONS present in media under the same conditions as those applied in OES and OAS analysis, and to demonstrate that the reactive species produced in the plasma are present in the liquid phase, and thus capable of reaching the cells. The media was exposed to the plasma discharge and incubated for 24 hours post treatment. Following incubation, Griess Reagent and Amplex™ Red reagent were used to quantify nitrite and hydrogen peroxide, respectively. As can been seen in figure 7, the plasma discharge produced species in a dose-dependent manner (P < 0.01 for nitrite and P < 0.001 for hydrogen peroxide), with no significant difference between the duty cycles at any doses of CAP tested. The highest concentration of nitrite found was just above 40 μM. For hydrogen peroxide the highest concentration was 13.8 μM.

**Figure 7.**
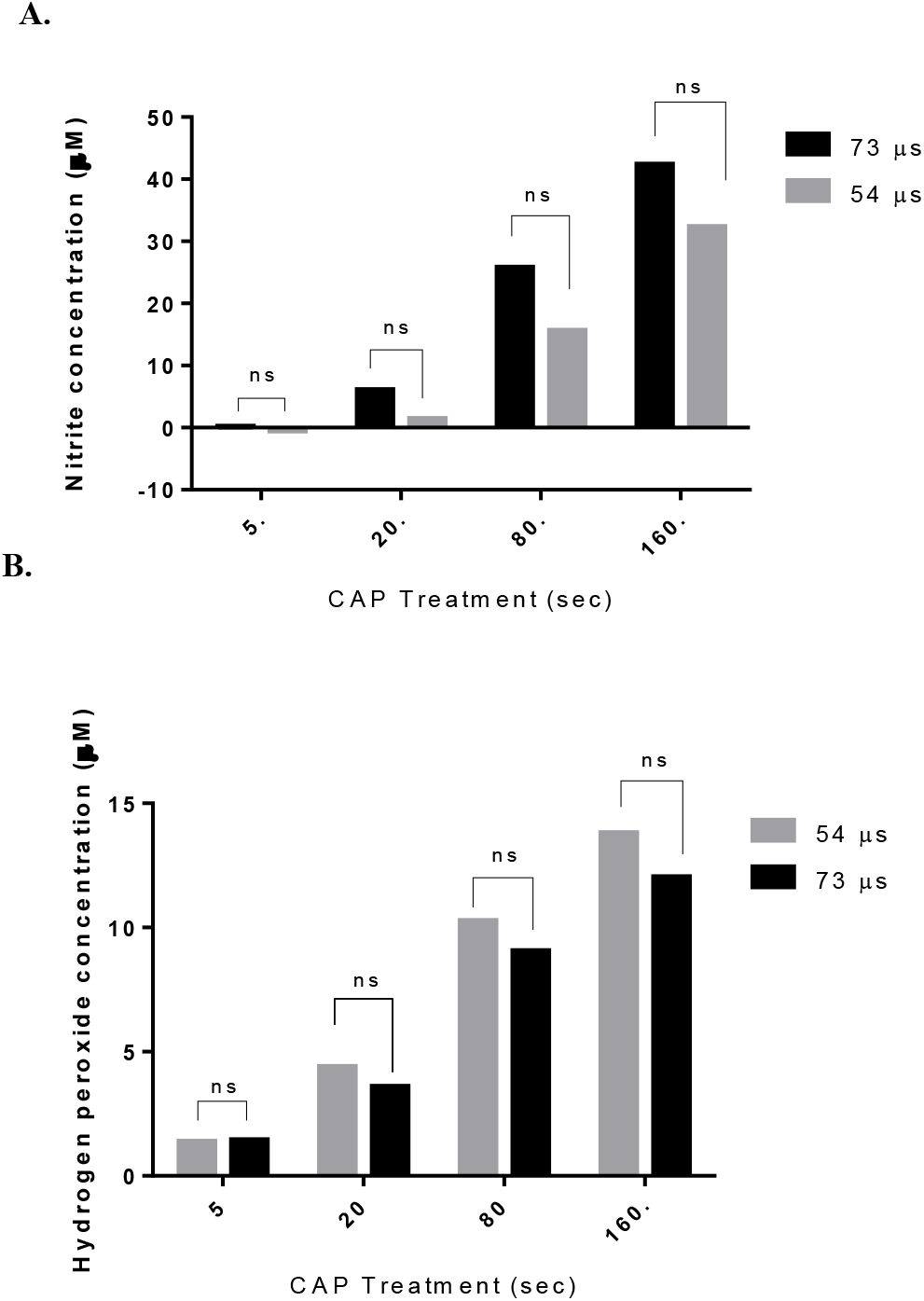
Concentrations of nitrite and hydrogen peroxide on media 24 hours post-treatment. DMEM without phenol red and in the absence of sodium pyruvate was used to quantify those RONS. Samples were exposure to four different doses of CAP (5, 20, 80 and 160 seconds). Plasma discharge was set as 240 V, 1000 Hz and either 73μs or 54μs. (A) Nitrite concentration was measured using Griess reagent. (B) Hydrogen peroxide concentration was quantified using Amplex™ Red reagent. Data was analysed using linear-regression interloping unknowns from standard curve. Interloped X values where then used to run a Two-way ANOVA with Sidak’s multiple comparisons test (ns P > 0.05). All experiments were performed in triplicate with a minimum of 4 replicates per experiment.

### ROS production in U-251 MG

After the analysis of RONS on plasma discharge (Figure 5 and 6) and in media (Figure 7), we wished to evaluate how the presence of those reactive species would be translated inside the cells. For this, cells were incubated with H2DCFDA, an indicator of reactive oxygen species in cells. Following treatment, cells were incubated for 3 hours, collected after running through CytoFlex Coulter and quantified using FL-1 green channel. Data was analysed using CytExpert software (Figure 8 A). Gate strategy used can be seen in supplementary figure II. The analysis of the dot plots shows evidence of ROS production at 160 seconds for both 73 μs and 54 μs (Figure 8 A). However, no statistically significant values were found (Figure 8 B).

**Figure 8.**
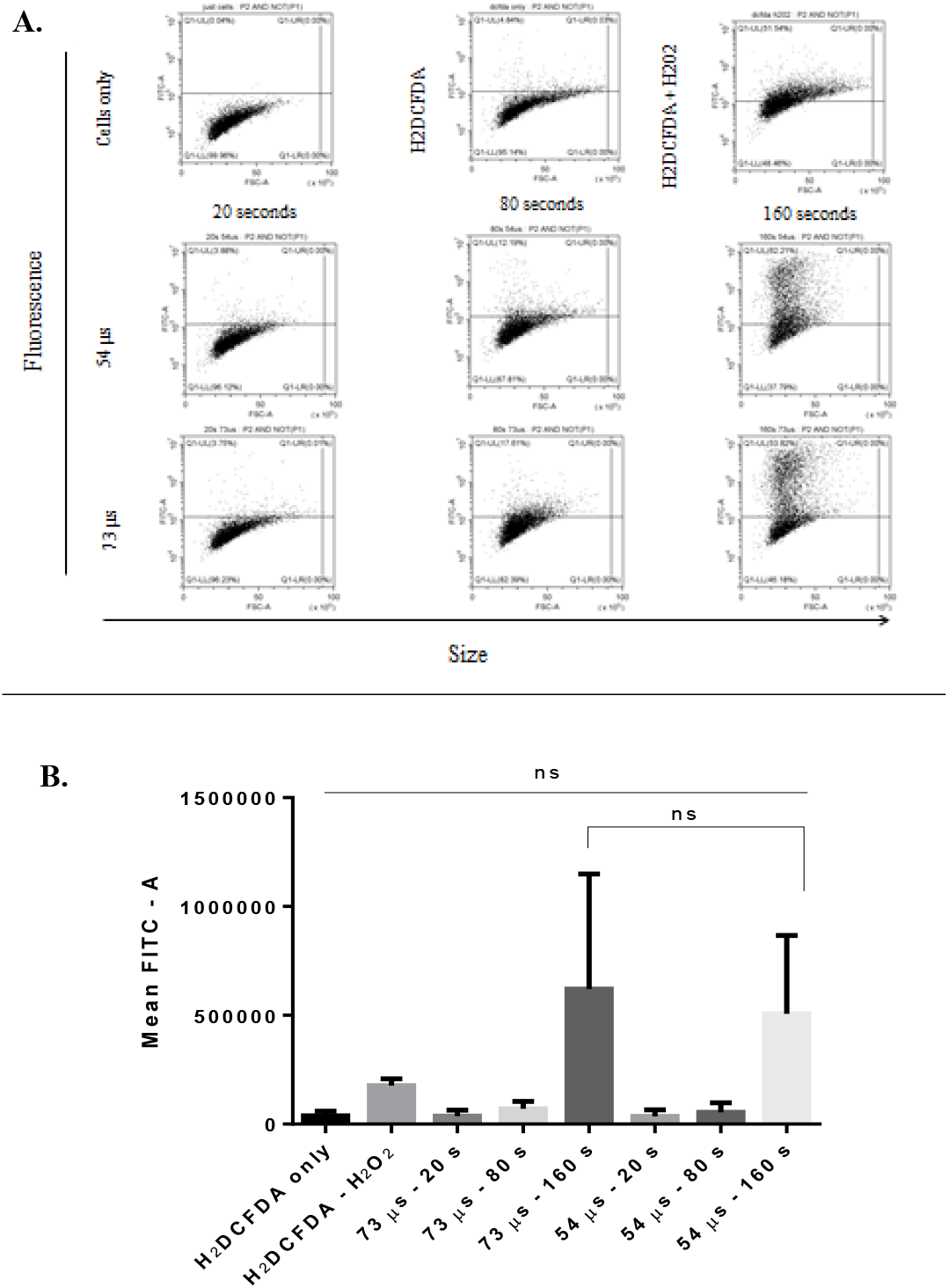
ROS production in U-251 MG. Cells were incubated with H2DCFDA and treated at three different doses of CAP using a plasma discharge of 240 V, 1000 Hz and 73 μs or 54 μs. (A) 3 hours post treatment cells were collected and analysed using CytExpert software. (B) The media of FITC channel was used to plot the values on columns. Two-way ANOVA with Sidàk multiple comparisons test was used (ns P > 0.05). Experiments were performed two independent times.

## Discussion

GBM is the most frequent tumour of the brain. Despite advances in science its five-year relative survival is only 6.8 % (Khani *et al.*, 2019; Ostrom *et al.*, 2019). This highlights the necessity for new therapeutic approaches. CAP induced cytotoxicity in cancer cells has been shown previously across a multitude of differing devices (Dubuc *et al.*, 2018). The pro-apoptotic effect of ROS in cancer cells has been demonstrated robustly in the literature and is largely what the current standard of care for GBM (radiotherapy combined with chemotherapy) is reliant on (Lo Dico *et al.*, 2019). A major disadvantage of these treatment options is the development of ROS resistant cells, indeed U-251 MG cells have been shown to be highly ROS resistant (Conway *et al.*, 2016).

We were able to assess the cytotoxicity effect of a pin-to-plate device, which works in a dose dependent manner. A higher cytotoxicity response was observed in cells after a longer incubated period post-treatment in contrast to an early cytotoxicity effect observed in the same U 251 MG cell line with the use of a di-electric barrier (DBD) system (Conway *et al.*, 2019). Propidium iodide (PI) uptake was also assessed and confirmed the delayed cytotoxicity effect of this pin-to-plate device. This indicates that it is most likely that a programmed cell death is occurring, rather than necrosis (Galluzi *et al.*, 2018). Previous diagnostic results (unpublished results) have reported that the pin-to-plate device has an optimum discharge frequency at 1000 Hz. We were able to assess the effect of 1000 Hz on GBM cell cytotoxicity, especially when DMEM in the absence of sodium pyruvate was used. Pyruvate is known to alters the outcome of treatment, due to its hydrogen peroxide scavenger properties and its activation of the Akt signalling pathway (Fernández-Gómez, *et al.*, 2006; Tornin *et al.*, 2019). The higher cytotoxicity observed in DMEM without sodium pyruvate indicates that hydrogen peroxide may also play a role. A stronger death response in the absence of pyruvate has previously been reported by Tornin and colleagues in osteosarcoma cells (Tornin *et al.*, 2019). The presence of sodium pyruvate appears to not have an effect in the first 24 hours post-treatment suggesting that different incubation times, may lead to different death pathways and/or ROS interaction. An important note is the absence of cytotoxicity when the pin-to-plate device is used without containment. Considering that the container box is used to entrap the RONS that have been generated by the plasma discharge, the removal of this container almost eliminates any gaseous species during treatment as they are quickly siphoned off via the laminar air hood that the entire device is set in. This is further evidence of RONS being the main contributor of the devices anti-cancer effect.

A ROS-dependent death was observed with the use of NAC. Difference in duty cycles produced no significant difference in cytotoxicity and NAC was able to reverse the cytotoxicity observed at both settings. There are several published reports of ROS-dependent death (Ahn *et al*, 2011; Vandamme *et al*, 2012; Siu *et al*, 2015; Bauer, 2019), interesting for this cell line, although with a different device, a ROS-independent mechanism had been reported (Conway *et al.*, 2016). Knowing the major role of RONS on cell viability and its dependence in cytotoxicity and the fact that pin-to-plate may lead to a ROS-dependent cell death, RONS was quantified in the plasma discharge, afterglow, media and intracellularly.

No NO or O was detected in the emission spectra obtained. Previous measurements of NO through the use of Draeger Tubes show low or no detectable emissions of NO, O, NOx (NO_2_, NO_3_, N_2_O_2_, N_2_O_3_, and N_2_O_4_), •OH and N_2_^+^ measured in air using OES for the DBD system. This is believed to be due high levels of ozone generated within the plasma discharge (Schmidt-Bleker *et al.*, 2015; He *et al.*, 2020), and the same explanation can be applied to the current study because high levels of ozone was observed. Fluctuations were observed over time between species as well as differences in the level of detected species over the sample area, possibly due to the dispersion of gas molecules throughout the system. Since the system is relatively static (no applied air flow), the highest energies should be seen in the centre of the system where the residence times of atoms and molecules allows them to stay in an excited state for longer. Therefore, when they begin to dissipate from the central portion of the system, where their population may build and saturates their relaxation process can cause emission events in the outer portions. The measured species are none-the-less measured at a relatively high level and at a close spread to one another. The main outlier for O_3_ measurement is from the same position (right from the centre) and duty cycle (73 μs) that yielded higher levels of N_2_, N_2_^+^, and OH, due to the fact that higher levels of those species leads to a higher number of events that contribute to reaction mechanisms that leads to O_3_ production. The central point also contains a higher level of O_3_ due to its longer residence time along with other reactive species. The differences observed, including between different duty cycles, can be regarded as negligible as it seems to have little impact on the treated cells.

The translation of the species into the media and intracellularly seems to follow the same pattern of the cytotoxicity effects and the discharge plasma RONS production, with no significant difference been observed between duty cycles. When RONS are present inside the cell it can induce apoptosis in gliomas and other tumours (Vandamme *et al,* 2012; Ahn *et al,* 2011; Siu *et al,* 2015; Bauer *et al.*, 2019). Evidence of higher intracellular ROS can be seen in figure 8 A with longer doses of CAP. We did not observe any increased intracellular reactive species using this method for shorter time points (20 s), although we do see evidence of hydrogen peroxide generation using Amplex Red and cytotoxicity, suggesting that intracellular antioxidants are effective at reducing RONS levels for shorter exposures to CAP, and that short lived reactive species may play an important role in the cytotoxic effects observed in GBM cells.

Hydrogen peroxide has been reported as the main anti-cancer reactive species involved in cell death using cold plasma (Yan, Sherman and Keidar, 2017) and its interaction with nitrite appears to be essential for biological effects in GBM cells (Girard *et al.*, 2016; Kurake *et al.*, 2016). Hydrogen peroxide and nitrite were quantified in a dose dependent manner. The presence of these two species gives an initial insight into the interaction of CAP/media for the pin-to-plate device. Although we quantified the presence of hydrogen peroxide, the concentration was low (up to 13.8 μM). According to Boehm *et al.*, 2018 an IC50 concentration of 315 μM of pure hydrogen peroxide and 185 μM of hydrogen peroxide on plasma active water was required to kill U251 MG cells. In the same study a dose response curve with concentrations of up to 1200 μM nitrite on the same cell line was performed, with nitrite *per se* not showing any cytotoxicity effect. The concentrations reported in this study were higher to the ones observed with the pin-to-plate device, indicating that hydrogen peroxide or nitrite are not the sole cytotoxicity factor. However, the low concentrations could be decayed products from short lived species, which may play a crucial role in cytotoxicity. In fact, others have reported that short exposure to direct CAP is sufficient to activate cancer cells into a sensitive state to ROS and RNS, including H2O2 and NO2^−^ (Yan *et al.*, 2018). Our results suggest that a similar process may also play a role in the enhanced cytotoxicity observed here underscoring the importance of direct exposure to CAP for cancer treatments.

## Conclusion

The results presented demonstrate that the pin-to-plate device successfully induces GBM cell death in a time dependent manner, and shows promise as a future therapy for glioblastoma multiforme. The approach offers a more diffuse type of filamentary discharge in air over those typically found with DBD designs which would be more suited to *in vivo* treatments. The device induced cytotoxicity appears to be dependent on ROS production. The quantification of hydrogen peroxide and nitrite in quantities less than those needed to produce cytotoxicity *per se* indicates that the interaction between those species and/or with others RONS may play a major role in sensitising cells to CAP. More studies will be performed to elucidate the role of hydrogen peroxide and its further generated species.

## Acknowledgements

This work is supported by Science Foundation Ireland Grant Numbers 14/IA/2626 (P.C., J.C.) 17/CDA/4653 (A.C., B.T., P.C., J.C.) and by the TU Dublin Fiosraigh Scholarship Programme (S.B., J.C.). The authors would also like to acknowledge funding from the Food Institutional Research Measure administered by the Department of Agriculture, Food & the Marine, Ireland (Grant number: 13F442) (C.S., P.C).

