## Supplementary Materials for "Pin Electrode Reactor: A novel cold atmospheric plasma device and its potential in glioblastoma treatment"

**Supplementary data**

Table I – Sidak's multiple comparisons test U372 MG

A) DMEM SP (-), 1000 Hz 24 hours PT / DMEM SP (+), 1000 Hz 24 hours PT

| Dose CAP (seconds) | Mean Diff. | 95% CI of diff. | Significant? | Summary |
| --- | --- | --- | --- | --- |
| 0. | -27.78 | -114.2 to 58.69 | No | ns |
| 5. | -119.5 | -206.0 to -33.05 | Yes | ** |
| 10. | -50.04 | -136.5 to 36.43 | No | ns |
| 20. | -48.00 | -134.5 to 38.47 | No | ns |
| 40. | -18.47 | -104.9 to 68.00 | No | ns |
| 80. | -34.62 | -121.1 to 51.85 | No | ns |
| 160. | -34.92 | -121.4 to 51.55 | No | ns |
| 320. | 5.224 | -81.24 to 91.69 | No | ns |

SP (-) = without sodium pyruvate; SP (+) = with sodium pyruvate; PT = post treatment

B) DMEM SP (-), 1000 Hz 96 hours PT / DMEM SP (+), 1000 Hz 96 hours PT

| Dose CAP (seconds) | Mean Diff. | 95% CI of diff. | Significant? | Summary |
| --- | --- | --- | --- | --- |
| 0. | 128.6 | 47.14 to 210.1 | Yes | *** |
| 5. | 536.6 | 455.1 to 618.1 | Yes | **** |
| 10. | 645.8 | 564.3 to 727.3 | Yes | **** |
| 20. | 529.5 | 448.0 to 611.0 | Yes | **** |
| 40. | 528.2 | 446.7 to 609.7 | Yes | **** |
| 80. | 419.9 | 338.4 to 501.4 | Yes | **** |
| 160. | 219.5 | 138.0 to 301.0 | Yes | **** |
| 320. | 61.28 | -20.20 to 142.8 | No | ns |

SP (-) = without sodium pyruvate; SP (+) = with sodium pyruvate; PT = post treatment

C) DMEM SP (-), 2500 Hz 96 hours PT / DMEM SP (+), 2500 Hz 96 hours PT

| Dose CAP (seconds) | Mean Diff. | 95% CI of diff. | Significant? | Summary |
| --- | --- | --- | --- | --- |
| 0. | -67.35 | -153.0 to 18.27 | No | ns |
| 5. | -89.42 | -175.0 to -3.804 | Yes | * |
| 10. | 110.2 | 24.60 to 195.8 | Yes | ** |
| 20. | -60.23 | -145.8 to 25.39 | No | ns |
| 40. | 12.95 | -72.67 to 98.57 | No | ns |
| 80. | 12.13 | -73.49 to 97.75 | No | ns |
| 160. | 7.553 | -78.07 to 93.17 | No | ns |
| 320. | 17.80 | -67.82 to 103.4 | No | ns |

SP (-) = without sodium pyruvate; SP (+) = with sodium pyruvate; PT = post treatment

D) DMEM SP (-), 2500 Hz 96 hours PT / DMEM SP (-), 1000 Hz 96 hours PT

| Dose CAP (seconds) | Mean Diff. | 95% CI of diff. | Significant? | Summary |
| --- | --- | --- | --- | --- |
| 0. | 289.1 | 207.6 to 370.7 | Yes | **** |
| 5. | 34.25 | -47.30 to 115.8 | No | ns |
| 10. | -49.97 | -131.5 to 31.59 | No | ns |
| 20. | -98.77 | -180.3 to -17.21 | Yes | ** |
| 40. | -4.900 | -86.45 to 76.65 | No | ns |
| 80. | 12.67 | -68.88 to 94.22 | No | ns |
| 160. | 22.39 | -59.17 to 103.9 | No | ns |
| 320. | 40.45 | -41.10 to 122.0 | No | ns |

SP (-) = without sodium pyruvate; SP (+) = with sodium pyruvate; PT = post treatment

E) DMEM SP (+), 2500 Hz 96 hours PT / DMEM SP (+), 1000 Hz 96 hours PT

| Dose CAP (seconds) | Mean Diff. | 95% CI of diff. | Significant? | Summary |
| --- | --- | --- | --- | --- |
| 0. | -485.1 | -570.6 to -399.5 | Yes | **** |
| 5. | -660.3 | -745.8 to -574.7 | Yes | **** |
| 10. | -485.6 | -571.1 to -400.0 | Yes | **** |
| 20. | -490.9 | -576.5 to -405.4 | Yes | **** |
| 40. | -510.3 | -595.9 to -424.8 | Yes | **** |
| 80. | -420.5 | -506.0 to -334.9 | Yes | **** |
| 160. | -234.3 | -319.9 to -148.8 | Yes | **** |
| 320. | -83.94 | -169.5 to 1.621 | No | ns |

SP (-) = without sodium pyruvate; SP (+) = with sodium pyruvate; PT = post treatment

Table II – Dose response curve

| Cell type | Conditions/parameters | IC_50_ | Hill slope | R^2^ |
| --- | --- | --- | --- | --- |
| U373 MG | DMEM SP (-) / 1000 Hz / 24 hours PT | 143.6 (116.5 ± 176.9) | -0.5293 | 0.3225 |
| U373 MG | DMEM SP (+) / 1000 Hz / 24 hours PT | 145.8 (119.1 ± 178.6) | -0.6714 | 0.3232 |
| U373 MG | DMEM SP (-) / 2500 Hz / 96 hours PT | 11.22 (10.09 ± 12.48 | -1.223 | 0.5489 |
| U373 MG | DMEM SP (+) / 2500 Hz / 96 hours PT | 12.58 (11.56 ± 13.68) | -1.290 | 0.6518 |
| U373 MG | DMEM SP (-) / 1000 Hz / 96 hours PT | 5.716 (5.308 ± 6.155) | -1.256 | 0.7039 |
| U373 MG | DMEM SP (+) / 1000 Hz / 96 hours PT | 36.38 (33.51 ± 39.48) | -1.035 | 0.7319 |

SP (-) = without sodium pyruvate; SP (+) = with sodium pyruvate; PT = post treatment; R^2^ = coefficient of determination.

**Figure I – Spectrum of plasma discharge**

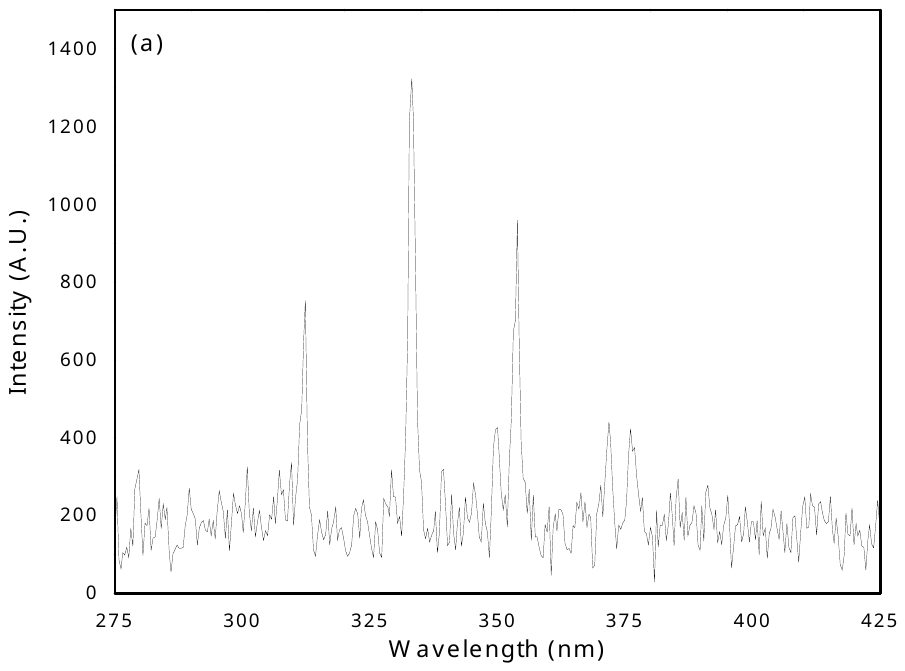

Figure 2 - **Spectrum obtained at a voltage setting of 240 V, duty cycle of 73 µs, and discharge frequency of 1000 Hz.** (A) Shows the spectral range in which OH and N_2_ of the SPS and FNS can be found. (B) The spectral range in which would find the 777 OI triplet.

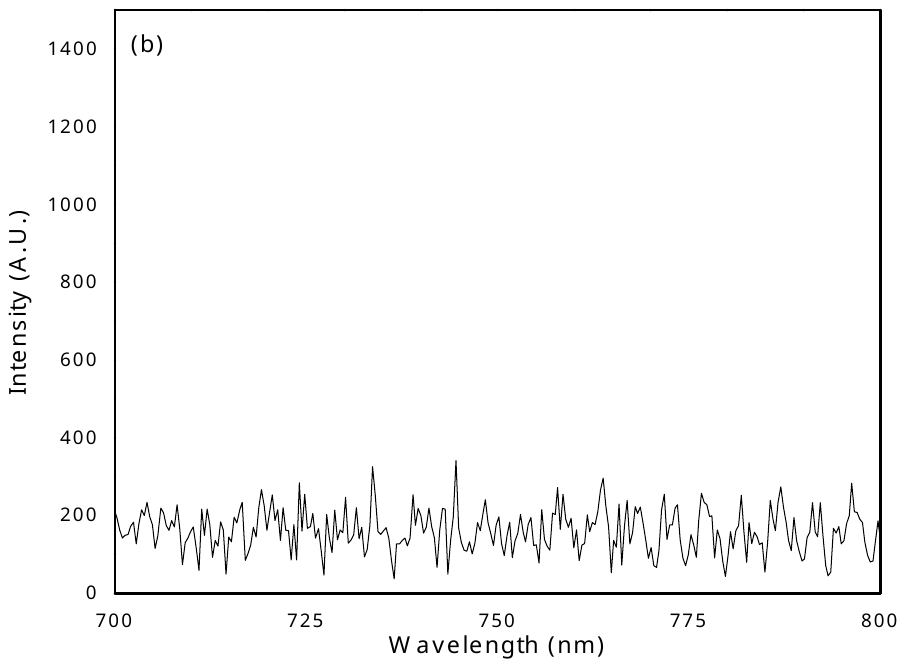

**Figure II - Gating strategy for H_2_DCFDA analysis**

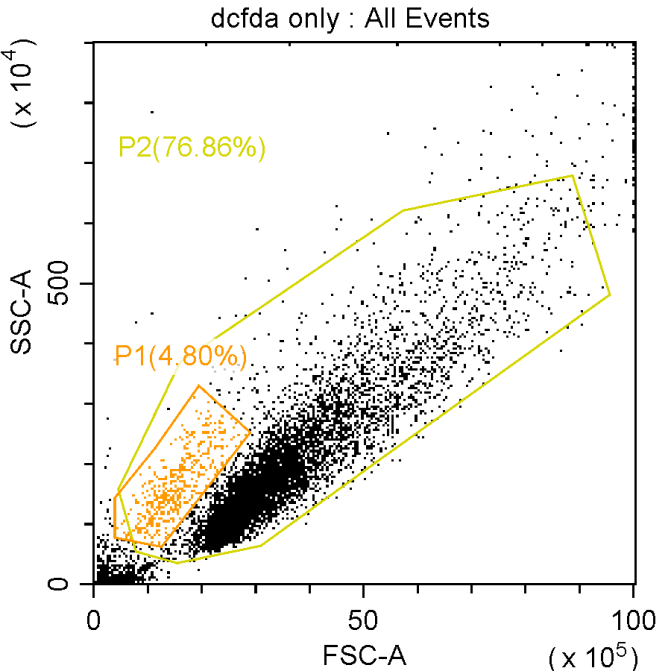

C.

B.

A.

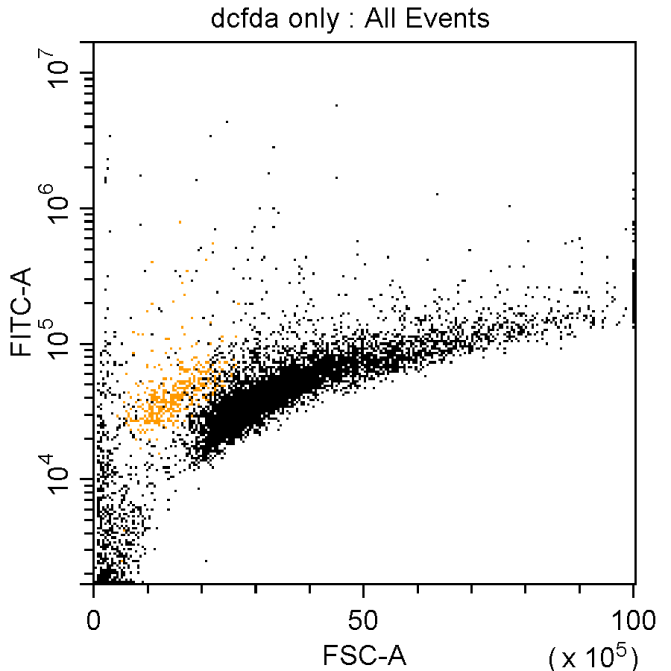

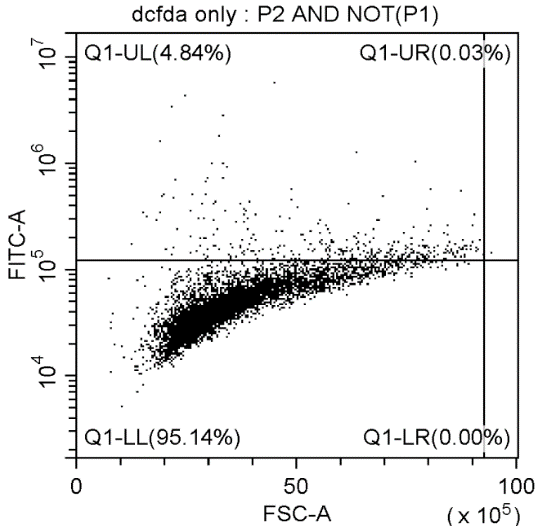

**Figure 3 – Gating strategy for H_2_DCFDA analysis.** A) Samples containing only H_2_DCFDA was plotted and a gate were draw around it (P2) eliminating debris, a second gate (P1) was also draw to be exclude of P2. B) Dot plot of all events C) Dot plot of P2 excluding P1.
